# Leave-One-Out-Cross-Validation Based Machine Learning Algorithms for Efficient, Scalable, and Robust Prediction of Artemisinin Resistance in *Plasmodium falciparum* Kelch13 mutants

**DOI:** 10.64898/2026.11.05.764763

**Authors:** Rajdeep Das, Smritikana Dutta, Souvik Bhattacharjee

## Abstract

**Background:** Emerging *Plasmodium falciparum* resistance to frontline artemisinin (ART) derivatives threatens global malaria control efforts. Current protocols assigning ART-sensitive (ART-S) or -resistant (ART-R) phenotypes to parasite strains with Kelch13 (K13) mutations remain time-intensive for real-time surveillance. Ring-survival assay (RSA), an *ex vivo* gold standard mandates laboratory-adaptation under culture conditions and frequently does not replicate clinical ART-responsiveness, thereby limiting its utility for rapidly triaging newly emerging K13 variants. In contrast, advent of artificial intelligence (AI)-derived models hold potential to screen >100 K13 variants within hours, enabling rapid near-real-time prediction of ART-S/R. Here, we engineered a panel of forty representative K13 variants by computing their relative deviations from wild-type K13 across ten physicochemical and structural descriptors, yielding a compact feature set which captures mutation-induced perturbations in folding, stability, and solvent exposure. These can be further utilized to train and validate AI classifier to predict resistance.

**Methods:** K13 wild-type and 39 mutants identified from clinical isolates were subjected to L1 regularised leave-one-out-cross-validation-based logistic regression on ten physicochemical and structural descriptors, namely propensity, isoelectric point (pI), GRAVY, aliphatic index (ALIN), instability index (II), hydrogen-bonding capacity (HB), steric hindrance (SH), solvent-accessible surface area (SASA), bond dissociation energy (BDE), and root-mean-square deviation (RMSD) features to classify them into ART-R/S, which was then followed by a linear regression with Lasso regularisation to predict their parasite clearance half-life (PC_1/2_) values. The two scripts were integrated and wrapped with Pyodide for web implementation.

**Results:** Linear regression model showed limited predictive power, yielding a maximum cross-validated Q^2^ of 0.110 with only ALIN. The L1-regularized logistic classifier instead showed robust performance efficiency (accuracy 80%, MCC 0.588, AUC 0.844) on ALIN, II and BDE.

**Conclusions:** While K13 mutations remain extensively characterized, a minimal, interpretable classifier based solely on physicochemical deviations from wild-type K13 is yet to be reported. Our findings indicate that such a classifier, based on physicochemical deviations of K13 mutants from the wild-type K13, can reliably predict ART-R phenotypes and would be valuable for prioritizing newly identified mutants for molecular surveillance and experimental validation. Training, and implementation scripts are available at: https://github.com/22rajdeep04/k13-ARP.git, and the web-tool can be accessed at: https://22rajdeep04.github.io/K13-ARP.

## INTRODUCTION

Malaria, mainly caused by *Plasmodium falciparum*, is a worldwide public health problem. According to WHO, an estimated 282 million global cases of malaria were recorded in 2024, with over 610,000 deaths (1). Although significant progress has been made during the last decade in reducing malaria-related morbidity and mortality, the concurrent emergence of drug resistance caused a major threat to worldwide malaria control efforts. Parasite resistance to well-known antimalarials such as chloroquine and sulphadoxine-pyrimethamine have already been reported, and rapidly spread over Asia and Africa (2,3). Artemisinin derivatives (ART) were subsequently introduced in 2004 as the frontline treatment for both acute and chronic forms of the disease (4). Originally isolated from the *Artemisia annua* in China (5–7), ARTs are widely recognized for their exceptional antimalarial potency. However, resistance to ART has since emerged in many regions after its first detection in South-east Asia (SEA) in 2008-2009 (8,9), and many more recently in eastern Africa (10–13). The poor pharmacokinetics of ART, especially its short half-life (<2 h), often leads to substantial failure in monotherapy, thereby necessitating its combination [as ART-based Combination Therapy (ACT)] with longer-lasting partner drugs such as lumefantrine, mefloquine and sulphadoxine–pyrimethamine (14,15).

Clinical emergence of ART resistance has primarily been attributed to single nucleotide polymorphisms (SNP) in the parasite’s *Kelch13* (K13; PlasmoDB ID: PF3D7_1343700) gene (16,17). *Plasmodium falciparum* strain 3D7 was first the clone derived from NF54 strain in Netherland, and is thought to be of African origin (18). It’s genetic stability and traceability, in addition to its drug susceptible background has proven to be an excellent system for ART-R studies (19,20). The K13 protein of 726 amino acids comprises of an *Apicomplexa*-specific N-terminus followed by three conserved domains, such as coiled-coil domain (CCD); a Broad-complex, tramtrack and bric-a-brac; (BTB)/the poxvirus and zinc-finger (POZ) domain and a C-terminal Kelch-repeat propeller (KRP) domain with six Kelch β-propeller motifs (21). Majority of mutations in K13 detected in SEA are limited to the KRP domains (22,23), although worldwide polymorphisms span the entire K13 sequence (24,25). Mutations such as K13 C580Y and R539T mostly dominant in SEA, and A578S in Africa (26,27). Other nonsynonymous K13 mutations associated with delayed parasite clearance in the Greater Mekong subregion (GMS) are F446I, Y493H, I543T, P553L, R561H, P574L and A675V (28,29). Recent reports have revealed the emergence of new ART-R K13 mutations in Africa, including P441L, C469F, C469Y, R561H and A675V (13,30,31). As of now, there are >200 K13 mutations identified in *P. falciparum* field isolates, of which the WHO classified 9 mutations as validated ART-R (F446I, N458Y, M476I, Y493H, R539T, I543T, P553L, R561H and C580Y) and 11 K13 mutations (P441L, G449A, C469F, A481V, P527H, N537I, G538V, V568G, P574L, F673I and A675V) as candidates with potential resistance. Non-statistically significant associated K13 mutations are D452E, C469Y, K479I, R515K, S522C, N537D, R575K, M579I, D584V, P667T and H719N) (1,32).

Ring-stage survival assay (RSA) established itself as a primary screen for the worldwide surveillance of ART-R. In RSA, clinical isolates are firstly adapted to tissue culture conditions, exposed to pharmacologically relevant dosage (700 nM) of ART for 6 h, washed free of ART and grown for an additional 66 h, and finally calculated for fast *versus* delayed clearance of infected erythrocytes in Giemsa-stained thin smears across the different fields of view, as compared to uninfected RBC. The results are expressed as percentage (33,34), where survival rate of ≥1% is generally classified as ART-R strain (http://www.wwarn.org/tools-resources/procedures/ring-stage-survivalassays-rsa-evaluate-vitro-and-ex-vivo-susceptibility). A clear correlation exists between the *in vitro* RSA and positive parasitemia following day 3 of ART treatment in patients, which further corroborated with mutations in K13. However, time-consuming RSA protocol suffers from several shortcomings, including low throughput screening, high variations in the exhaustive evaluation of microscopic readout, and poor correlation with *in vivo* resistance seen in clinical samples post-ART treatment. Further, visual inspection to distinguish developmentally viable and arrested parasites *versus* the dead pyknotic forms in thin blood smears often leads to assay variability (35,36). Other variations in the RSA protocol (37–39) and flow cytometry-based counting of viable parasites (35,36,40), have since been implemented and are comprehensively validated. A recently reports extended recovery RSA and quantified *P. falciparum* DNA in treated *versus* untreated control cultures by quantitative PCR (qPCR) under extended incubation period using known parasite clearance half-life values (PC_1/2_), which permitted comparative assessment of multiple samples and eliminated variability (35). Nonetheless, adaptation of *in vivo* field isolates directly from patients, stored subsequently *ex vivo*, and finally expanded for *in vitro* culture conditions can often be challenging and requires expertise in technical handling. Most of the *ex vivo* stored *P. falciparum* field isolates with K13 mutations either fail to revive or do not adapt to the *in vitro* culture conditions.

Artificial intelligence (AI) presents an attractive potential to strengthen malaria control efforts and predict future endemic outcomes (41–43). AI tools adopt machine learning and neural networks algorithms to process complex large-scale data and provide a comprehensive understanding of the dynamics of malaria transmission (43–45). AI-powered platforms have also shown potential in minimizing manual data entries as well as improve the efficacy of prediction of disease pattern (46). In marked contrast to the limitations of RSA and its advances in detecting parasites in blood smears, AI-driven tools can more accurately identify parasites, especially those with the rapid emerging newly-identified variants of K13 mutants in Africa (47,48). AI models, once trained with large previously published genomic datasets can identify these variants which occur at low frequency across the populations. A recently launched Malaria DREAM challenge, based on previously published datasets of transcriptome profiles in *P. falciparum isolates* from >1,000 patients (49,50), predicted ART-R by blindly evaluating algorithms to predict the *in vitro* drug response of independent isolates with expression data prior to and after exposure with ART derivatives (51). Prior studies have attempted to predict ART-R using machine learning models like Light Gradient Boosting Machine (LightGBM), eXtreme Gradient Boosting (XGBoost), Random Forest and Gaussian Process Regression, trained on gene expression level, which showed a maximum Area Under the Receiver Operating Characteristic curve (AUROC) of 0.86 (52–55). These models require transcriptomic data to predict ART-R, which could be challenging in impoverished regions.

In this study, we subjected the forty K13 mutants through ten distinct physicochemical and structural descriptors, thereby to train a logistic regression and linear regression model to predict the resistance and parasite-clearance half-life (PC_1/2_) of emerging strains with K13 mutations. ART-R due to K13 mutations is a complex and compound phenotype. Thus, the use of specific physicochemical parameters such as propensity, isoelectric point (pI), grand average of hydropathicity (GRAVY), aliphatic index (ALIN), instability index (II), bond dissociation energy (BDE) and total hydrogen bonds (HB), and structural descriptor such as solvent accessible surface area (SASA), steric hindrance (SH) and root mean square deviation (RMSD) allowed us to evaluate the resistance of the emerging strains with mutations in K13 holistically.

## METHODS

### Data Collection

We compiled a list of K13 mutants across different studies (21,32,56–58) to train our models. The list was supplemented with more mutants from large scale surveys done in Africa (56,59,60) and meta-analysis of the WWARN database (61,62). Finally, a composite list was prepared with forty variants in K13, including the ART-S K13 wild-type protein. The *P. falciparum* 3D7 was considered as the reference and we performed all analyses on the protein sequence accessed from UniProt (https://www.uniprot.org/) with accession code Q8IDQ2_PLAF7, identified from the whole genome sequencing of *P. falciparum* 3D7 (https://plasmodb.org/plasmo/). The composite **Table 1** enlists the mutations sampled for the study, along with degree of resistance and their PC_1/2_ values. We sampled mutations from the three most malaria endemic regions of the world, Sub-Saharan Africa, South East Asia and South Asia. **Figure 1** shows the global distribution of the K13 mutations identified across these endemic regions.

**Figure 1.**
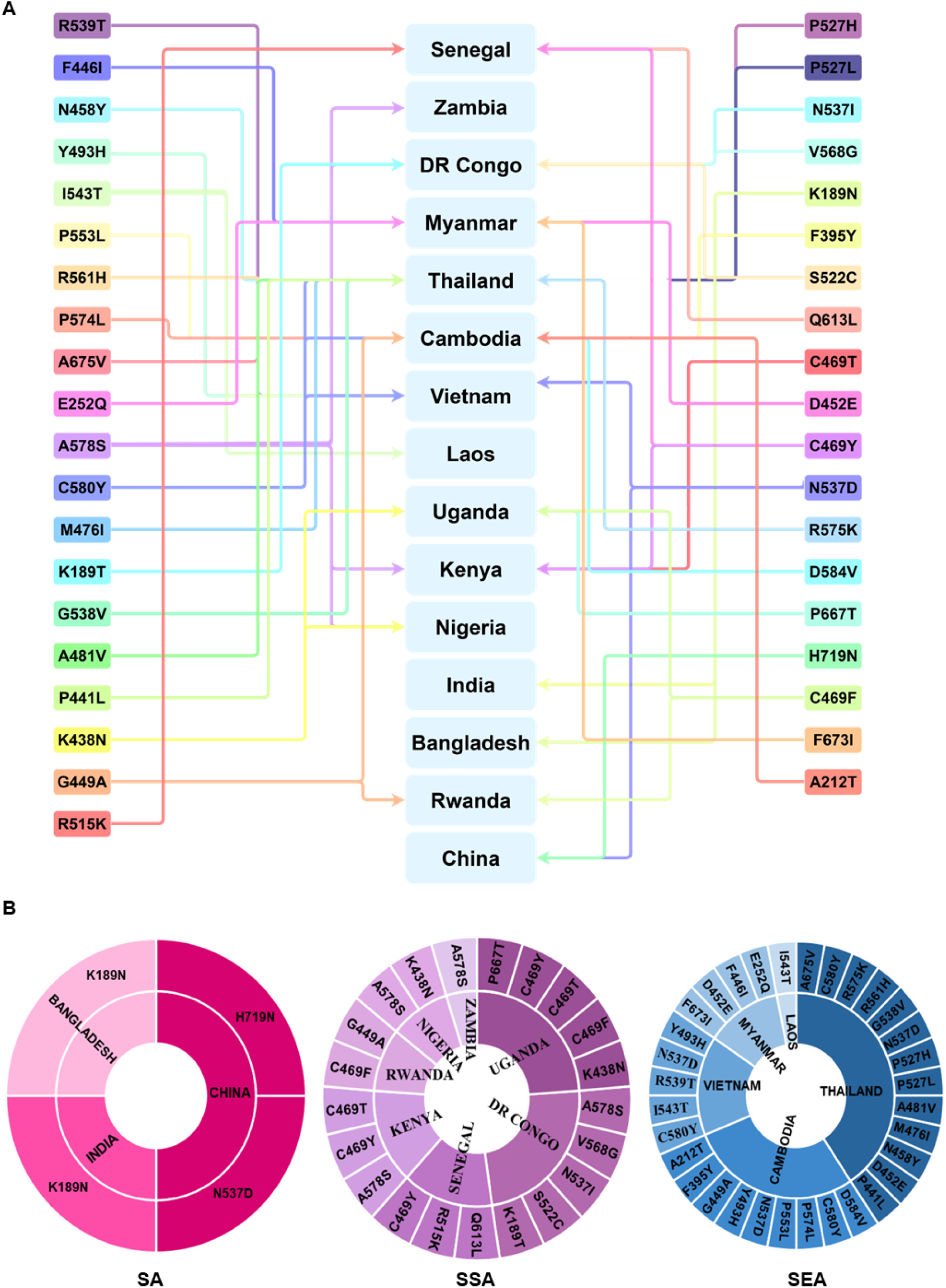
Global distribution of mutations in *Plasmodium falciparum* K13 protein across different countries in South Asia (SA), South-East Asia (SEA) and Sub-Saharan Africa (SSA). **A.** Schematic map showing the prevalence of K13 mutations detected in malaria surveillance of endemic regions in SA, SEA and SSA. **B.** Circle graph indicating sub-continent-wise presence of emerging and WHO-recognized K13 mutations in the different geographical malaria-endemic niches across the globe which serve as breeding grounds for the development of new drug-resistant K13 variants in *Plasmodium falciparum*.

**Table 1.**
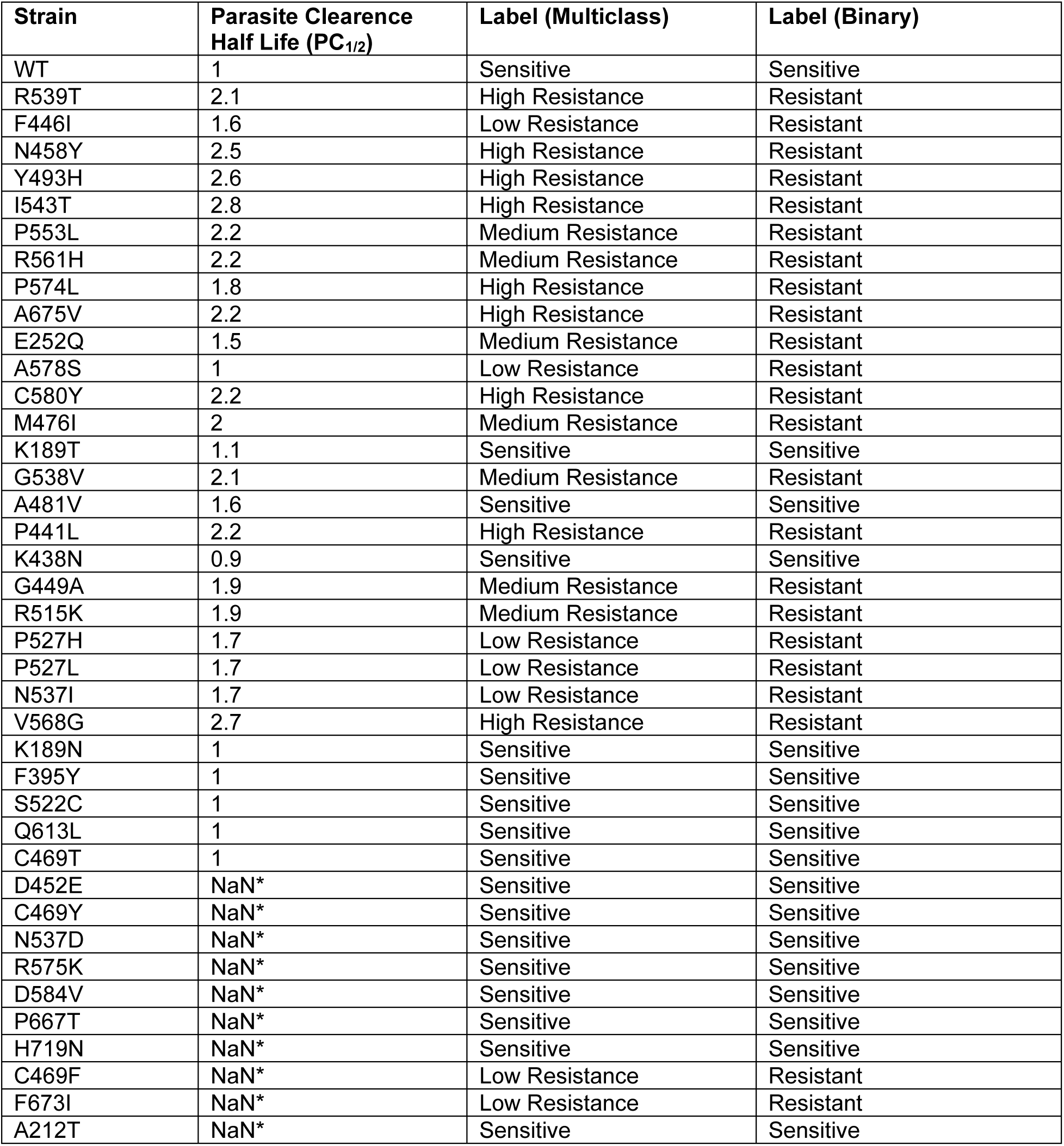
Resistance profiles, parasite clearance half-life and resistance labels of K13 mutants considered during the study. Linear regression model was trained on the log transformed parasite clearance half-life as the target variable (n=30), while the logistic classifier was trained on the discrete binary labels (n = 40). The values were corroborated by biochemical and cell viability assays (RSA) and clinical drug resistance profiles of the K13 variants. Since some of the mutants are yet to be characterised by RSA, their parasite clearance half-life remains unknown (Not a Number, NaN). Parasite clearance half-life has not been determined for the corresponding variants and were not included in the linear regression training and validation set.

| **Strain** | **Parasite Clearence Half Life (PC<sub>1/2</sub>)** | **Label (Multiclass)** | **Label (Binary)** |
|---|---|---|---|
| WT | 1 | Sensitive | Sensitive |
| R539T | 2.1 | High Resistance | Resistant |
| F446I | 1.6 | Low Resistance | Resistant |
| N458Y | 2.5 | High Resistance | Resistant |
| Y493H | 2.6 | High Resistance | Resistant |
| I543T | 2.8 | High Resistance | Resistant |
| P553L | 2.2 | Medium Resistance | Resistant |
| R561H | 2.2 | Medium Resistance | Resistant |
| P574L | 1.8 | High Resistance | Resistant |
| A675V | 2.2 | High Resistance | Resistant |
| E252Q | 1.5 | Medium Resistance | Resistant |
| A578S | 1 | Low Resistance | Resistant |
| C580Y | 2.2 | High Resistance | Resistant |
| M476I | 2 | Medium Resistance | Resistant |
| K189T | 1.1 | Sensitive | Sensitive |
| G538V | 2.1 | Medium Resistance | Resistant |
| A481V | 1.6 | Sensitive | Sensitive |
| P441L | 2.2 | High Resistance | Resistant |
| K438N | 0.9 | Sensitive | Sensitive |
| G449A | 1.9 | Medium Resistance | Resistant |
| R515K | 1.9 | Medium Resistance | Resistant |
| P527H | 1.7 | Low Resistance | Resistant |
| P527L | 1.7 | Low Resistance | Resistant |
| N537I | 1.7 | Low Resistance | Resistant |
| V568G | 2.7 | High Resistance | Resistant |
| K189N | 1 | Sensitive | Sensitive |
| F395Y | 1 | Sensitive | Sensitive |
| S522C | 1 | Sensitive | Sensitive |
| Q613L | 1 | Sensitive | Sensitive |
| C469T | 1 | Sensitive | Sensitive |
| D452E | NaN* | Sensitive | Sensitive |
| C469Y | NaN* | Sensitive | Sensitive |
| N537D | NaN* | Sensitive | Sensitive |
| R575K | NaN* | Sensitive | Sensitive |
| D584V | NaN* | Sensitive | Sensitive |
| P667T | NaN* | Sensitive | Sensitive |
| H719N | NaN* | Sensitive | Sensitive |
| C469F | NaN* | Low Resistance | Resistant |
| F673I | NaN* | Low Resistance | Resistant |
| A212T | NaN* | Sensitive | Sensitive |

### Propensity Calculator

We built an accurate propensity calculator which not only measured the propensity estimates with likelihood that a specific amino acid sequence will form a particular structural feature, physical property, or behaviour (63) but also presented an output for the K13 mutant sequence. The propensity was calculated using Chou-Fasman’s propensity table for amino acids, by first running a sliding window algorithm (64). The average ‘alpha, beta and turn propensities’ were calculated and loaded into a ‘pandas DataFrame’ (https://pandas.pydata.org/pandas-docs/stable/reference/api/pandas.DataFrame.html), which returned only 5 windows wherein the mutation affected the sequence and provided the output as a ‘propensity_comparison_{input mutant}.csv’ file. The python script for the calculator is publicly available in the GitHub repository (https://github.com/22rajdeep04/k13-ARP.git).

### Isoelectric Point Calculator 2

The IPC2.0 calculator was used to gain molecular insights into the isoelectric point (pI) at the local environment of the mutation (https://ipc2.mimuw.edu.pl/); (65). IPC2.0 runs on an advanced machine learning algorithm that was trained on over a 100,000 peptides and proteins using convolutional neural networks and support vector regression for peptides and proteins, respectively, that can predict the pI of the input protein sequences. The amino acid windows generated by the propensity calculator from each mutation was parsed into IPC2.0 to obtain the predicted pI value of the local environment around the site of substitution.

### Multiple Protein Profiler

We uploaded the same windows into Multiple Protein Profiler 1.0 (https://mproteinprofiler.microbiologyandimmunology.dal.ca/) (66) to calculate the GRAVY, aliphatic index (ALIN) and instability index (II). Although it also calculated other parameters such as charge and molecular weight; they were discarded since they were outside the purview of the current study, and only the three parameters were used for developing the algorithms, which could accurately predict the physicochemical parameters with extreme precision iteratively of multiple sequences at once.

### AlphaFold

The full-length mutant sequences of K13 received from the Propensity calculator were uploaded onto the Google DeepMind’s AlphaFold 3.0 server to obtain the predicted 3D Structure (https://alphafoldserver.com/) (67). AlphaFold 3.0 outperformed all its predecessors and the benchmark in terms of prediction accuracy and structure optimization in context to proteins and DNA. We further analysed the structure of the mutant proteins using **ChimeraX v1.12** (https://www.cgl.ucsf.edu/chimerax/) (68,69), a next-generation molecular visualization and analysis software which allowed us to calculate the total number of hydrogen bonds (HB), and the clash score. The difference in clash score of the mutant sequence from the wild-type K13 was used as a metric for measuring steric hinderance (SH). From the literature, we observed minor differences between the number of new HB formed in high resistance strains and low resistance strains. We aligned the predicted 3D structures of the mutants over the WT model on Schrodinger’s **PyMol** (https://www.pymol.org/) (70) to find the Root Mean Square Difference or RMSD.

### NetSurfP3.0

We uploaded the fasta sequences of the mutants on the NetSurfP3.0 server (https://bio.tools/netsurfp-3.0/) (71), that uses advanced machine learning algorithm to provide the relative and absolute surface area or solvent accessible surface area (SASA). To find the solvent accessible surface area or SASA, we considered the absolute difference between the K13 amino acid in wild-type from the mutant protein. The model runs a convolutional neural network and can predict the surface area with extreme precision. Additionally, it is also considerably faster than the prior version and other similar models.

### PEPCONF-DB

We accessed the PEPCONF Database (72), from the following link: ‘https://github.com/aoterodelaroza/pepconf.’ The database was composed of conformational energies of dipeptides, tripeptides, disulfide bond-rich proteins and many bioactive compounds. We first iterated over the amino acid sequence to find residues on either side of the site of mutation, and calculated the difference between the BDE of respective residues and found the absolute mean for our input data. The python script for the same can be accessed through the provided GitHub repository under “bde_deviation.py.” The programme performs the necessary calculations and returns the core BDE deviations on the terminal.

### Data Pre-processing and feature selection

The numerical values from each tool were selected and the absolute deviation between the mutant and the wild-type K13 sequence was measured. Since the local parameters were passed through the servers as five separate amino acid sequences, we considered the mean absolute deviation of the mutants from the wild-type K13 for those, but only the absolute deviation for global parameters such as steric hindrance. These values were then stored along with the labels obtained from prior studies in a ‘K13_complete.csv’ file, that can be found in **Supplementary table 1**.

To reduce the latency and the need for extensive data frame slicing to work on the appropriate data with individual models, two distinct files were created with only the pertinent labels in each namely, ‘K13_binary’ and ‘K13_pc’, wherein K13_pc contained continuous data of the PC_1/2_ ratios of K13 mutants to wild-type K13 and K13_binary contained the resistance labels. Respective files are shown as **Supplementary table 2** and **Supplementary table 3**, respectively. **Figure 2** displays the sequentially adopted data pre-processing and algorithms applied to build and deploy the models.

**Figure 2.**
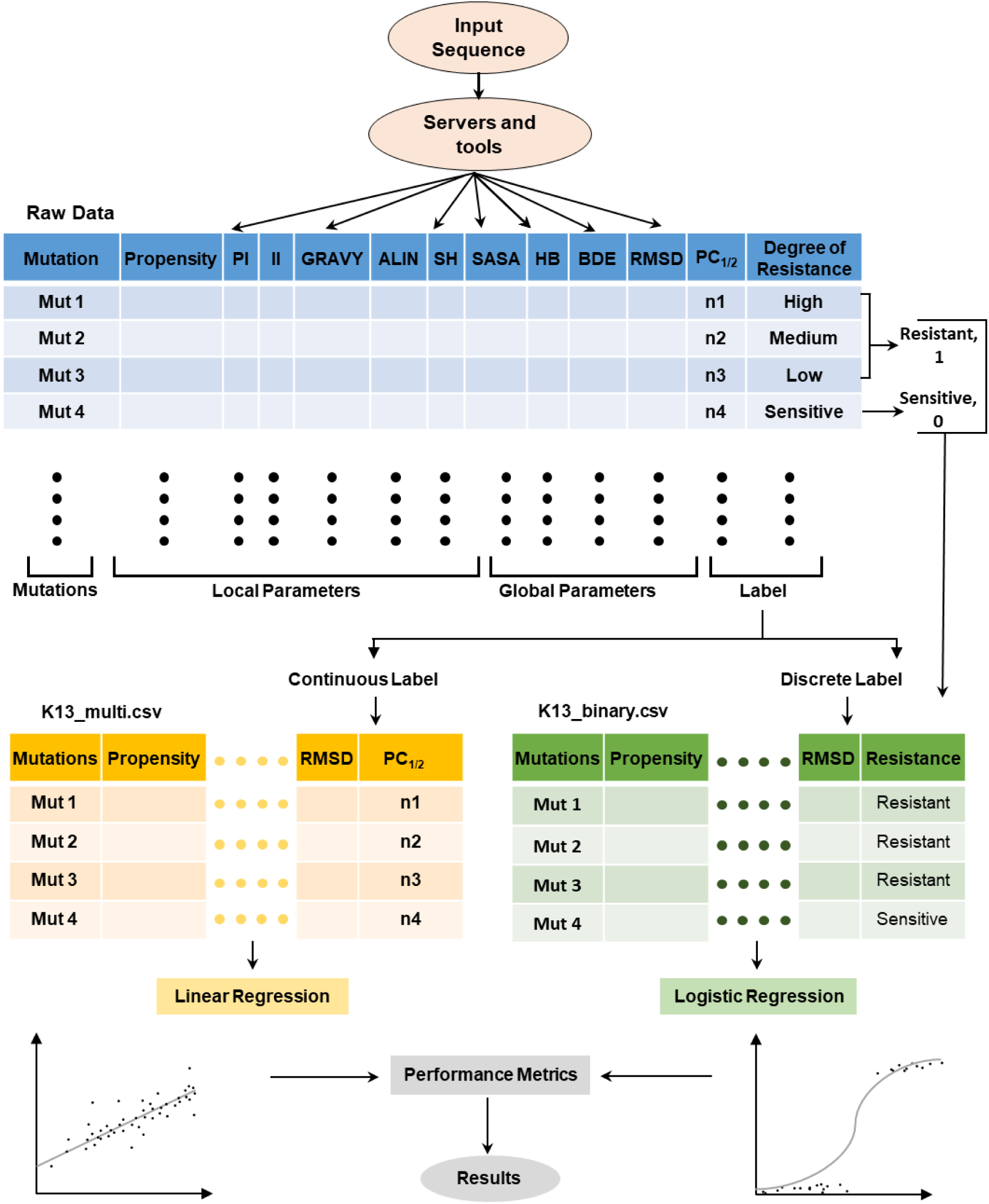
Systematic workflow diagram for robust data pre-processing and machine learning processes followed during this study. Individual K13 mutants, parsed by high-throughput servers such as Multiple Protein Profiler and developed scripts, to quantify the deviation of K13 mutants from wild-type for an array of 10 physicochemical and structural descriptors. Labels were segregated for computational parsimony and applied for feature selection and regression pipelines to datasets, as appropriate. A L1 regularised linear regression was applied to predict the parasite clearance half-life and an L2 regularised logistic resistance to predict the discrete probability of resistance [P(Resistance)]. The models were further analyzed using various tests and results compiled.

We then subjected the labelled data for the two supervised learning algorithms, namely logistic regression and linear regression. We used the Scikit-Learn Library in python (https://scikit-learn.org) to build and deploy the models (73). Before making the final fit model, the parameters that were most relevant biologically and computationally were selected. From the list with 10 initial features, 3 parameters for logistic regression and 1 parameter for linear regression were short-listed after an exhaustive and intensive feature selection pipeline over the dataset. We employed a Nested Leave-One-Out Cross-Validation (Nested LOOCV) method for the same, where for each training fold different features were tested and only the top k-most significant signals were fitted for the deployed models (74).

### Machine Learning

The features with z-score normalisation were standardised within each fold using the “StandardScaler” function of the sci-kit learn library (73). Additionally, considering the small size of the dataset, we used LOOCV or Leave-One-Out Cross Validation for training and testing the models. LOOCV holds on to one sample for testing and trains the model on the rest of the sample, *i.e.*, for the chosen set (n=40), the model leaves one sample and trains on 39 sample, then tests the left-out sample, following which it loops over the all samples sequentially. The process maximises the training data available at each fold, furthermore, the nested standardisation and feature selection prevented data leakage. For small datasets, LOOCV and Nested LOOCV generates the most optimal outcome in terms of prediction accuracy for supervised learning algorithms like Logistic Regression and Linear Regression. LOOCV provided an approximately unbiased estimation of generalisation performance and allowing overfitting to be detected than being masked. The strength of regularisation for both models was selected using a 5-fold CV over a fine log-spaced grid to ensure stable resolution of feature selection boundaries(75) using the NumPy library (https://numpy.org/) (76).

### Logistic Regression

We applied a L1 Penalty for embedded feature selection due to the limited size of the dataset and class weighting to mitigate the class imbalance. We calculated the selection frequency of all ten features across 40 folds, representing individual variants. and only retained the three features out of those selected in 100% of the folds as the stable core feature set, with ALIN, BDE and II (**Figure 3A-B**), corroborated by an accuracy *versus* k-sweep confirming peak performance in terms of accuracy and Matthews Correlation Coefficient (MCC) at k=3 (**Figure 3C**). Although SASA and Propensity also showed 100% selection rates, we only considered ALIN, II and BDE as the final feature set since they showed the highest |r| with resistance (**Figure 3D**). The decision threshold was further lowered to 0.4, that significantly reduced the number of false negative predictions made by the model from 7 to only 3, and consequentially it also increased the accuracy of the model from 76% to 80% (**Figure 3C**). This step was deemed crucial since false negative predictions (ART-R mutants being labelled as ART-S) would have significantly impacted the patient survival rates. For post feature selection, we refit the final deployed model on the complete dataset using the three core features with L2-regularised logistic regression and class weighting. To assess multicollinearity among the surviving features for logistic regression, we checked the Variance Inflation Factor (77) (**Figure 3E**).

**Figure 3.**
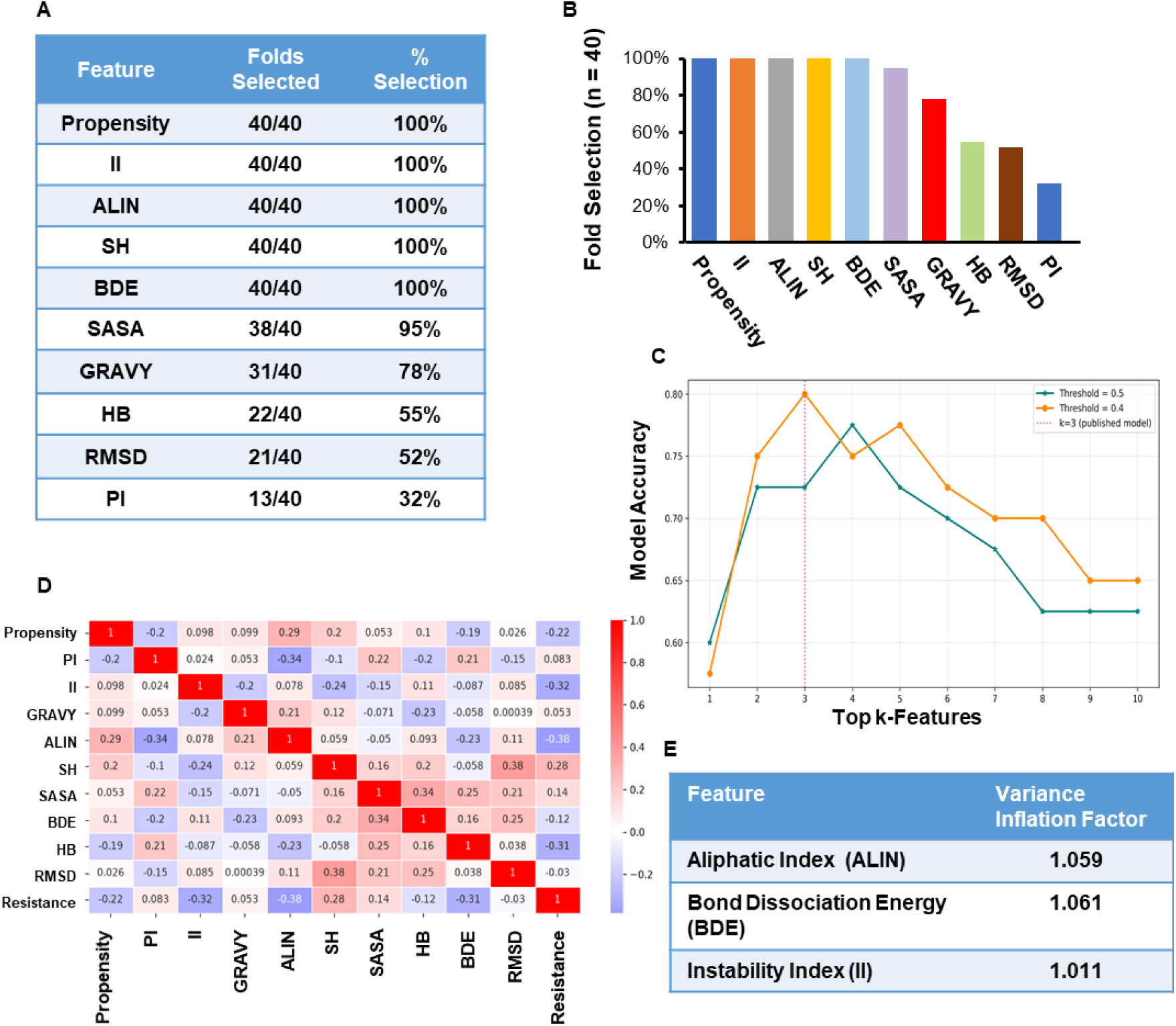
Basis for the selection of final set of features used in the logistic regression model. **A-B.** Table (A) and graph (B) showing percentage of inner cross-validation selection rates for the 10 individual features among the 40 selected K13 variants. Fine-spaced grid search identified the most informative features relevant to the model. Solvent-accessible surface area (SASA), grand average of hydropathy/ hydropathicity (GRAVY), hydrogen-bonding capacity (HB), root-mean-square deviation (RMSD) and isoelectric point (PI) parameters revealed <100% selection frequencies. Hundred percent selection frequencies of propensity, instability index (II), aliphatic index (ALIN), steric hinderance (SH) and bond dissociation energy (BDE) helped narrow core feature set to only 5 parameters. **C.** Accuracy *versus* k-sweep graph indicating that the model reached peak performance at k = 3, thereby finally training the model only on 3 features, namely ALIN, II and BDE for best accuracy. **D.** Label encoded heatmap showing the correlation between the features and the reported resistance, thus confirming the use of ALIN, II and BDE as core features. The Pearson’s correlation, |r|, with resistance was highest for ALIN (0.38), followed by II (0.32), and BDE (0.31). **E.** Table showing the variance influence factor (VIF) check and revealing ALIN, II and BDE showed no multicollinearity as VIF scores were between 1-1.1.

### Linear Regression

To identify the features that are most associated with the target, we performed an embedded L1-based feature selection using LassoCV (https://scikit-learn.org/stable/modules/generated/sklearn.linear_model.LassoCV.html). We used the Nested LOOCV framework mentioned above to find stable selection rates of the parameters. Although SH was selected in 3% of the folds (**Figure 5A**), it was hardly distinguishable from noise. Hence, the only parameter used for the model was ALIN, that was selected in 76% of the folds (**Figure 5A-B**). Given the log-linear relationship between clearance half-life and the clearance rate constant, the target was log transformed. It empirically improved the relationship, increasing LOOCV Q^2^ from 0.066 to 0.110 (**Figure 5C**). To stress test our model, we performed a permutation test (**Figure 5D**) on the model over 1000 target label shuffling’s. For future use, the final ALIN-only linear regression model was refit on the complete dataset.

### Model Integration and Web Implementation

In the training script, we integrated a configuration file generation block that stored the necessary datapoints like feature names, means and standard deviations for each feature to be used for the standardisation step, along with the coefficients and intercept as ‘classifier_config.json’ and ‘regression_config.json’ for the logistic and linear regression respectively. The configuration file also stored the decision threshold of the logistic regression, *i.e*., 0.4. Thus, the integrated approach ensured that the configuration file was regenerated with every training run.

We adapted the sliding window logic from the propensity calculator script and used the same to generate the windows required for calculating ALIN and II. From Multiple Protein Profiler (78), we imported the ALIN and II functions, which calculated the model acceptable parameters that, along with the calculated BDE from the aforementioned script, were passed through mean scaling using parameters stored in the configuration files. These standardised and scaled values were used for further calculation of the linear predictor (z) which was then used for calculating the probability of resistance by the logistic regression and the parasite clearance half-life (PC_1/2_) by the linear regression model using the data stored in the configuration files. The final script can be accessed through ‘main.py’ under the GitHub repository of the project.

To increase the usability of the programme, we wrapped the programme as a web tool using Pyodide that allows pythons script to run directly on the web. The client-side execution will allow the end user to enter the mutation to predict the resistance profile without the need to download any scripts or python packages, which significantly increased the accessibility and privacy, since the mutations were never uploaded to any servers. The tool first checked if the resistance profile of the input mutation is known and returned the parasite clearance half-life and resistance. In cases where the input strain was not characterised, it predicted the same using the stored model parameters. The web tool can be accessed through: https://22rajdeep04.github.io/K13-ARP.

### Testing novel and emerging mutants in K13

We identified 18 newly discovered mutant strains of the parasite in SA, SEA and SSA (79–81), whose resistance profile were yet to be determined and the RSA data was unavailable. We used the web tool to predict the resistance of these novel mutants and cross-validated using the models directly. **Supplementary table 4** shows the resistance profile and the parasite clearance half-life of the uncharacterised mutant strains, as predicted by the models.

## RESULTS

### Dataset for physicochemical and structural evaluation between K13 wild-type and mutants

Our curated dataset comprised of 40 variants of *P. falciparum* K13 with 10 physicochemical descriptors or features. We observed that 16 of the selected K13 variants were sensitive to ART while 23 were ART-R. We used the curated dataset to train and deploy two machine learning algorithms, one to classify and predict resistance in *P. falciparum* and one to predict the more clinically informative PC_1/2_.

### Logistic regression model achieves high accuracy in resistance classification

Using the above dataset, we employed the nested-LOOCV method for feature selection. Results, as shown in **Figures 3A-B**, revealed the selection rates of individual features in each inner LOOCV fold. We observed that Propensity, BDE, SH, ALIN, II and Propensity were highlighted in 100% of the selection folds. In nested LOOCV, the outer folds iterated over n samples, where n = 40, leaving one out for testing, while the inner LOOCV fold iterated over the remaining n-1 samples and selected the features and tune hyperparameters (n-1= 39). Nested LOOCV prevented the selection bias from overlapping features, consistent with established methods for small sample machine learning model (82,83). To further confirm the number of features that would likely yield the most optimal results, we plotted accuracy against the number of features (k). As shown in **Figure 3C**, we observed that the model reached 80% validation accuracy when only three features (BDE, ALIN and II) were selected. However, the accuracy dropped to 76% upon addition of SH and SASA as core features. The heatmap generated in **Figure 3E** indicated relatively low correlation between the target and any individual features, which suggested that any singular feature cannot completely capture the resistance phenotype of the parasite. Hence, a complex input was necessary to accuracy predict the same. **Figure 3D** also depicted that AI, II and BDE have the highest correlation (|r|) with resistance (0.38, 0.32 and 0.31 respectively). To rule out multicollinearity in our selected features, we calculated the VIF, or Variance Inflation Factor (77). **Figure 3E** indicated that the VIF values for all three features were between 1 and 1.1, thereby leading us to conclude that only AI, II and BDE should be used as true descriptor for the logistic regression model.

After selecting the final feature set, we applied LOOCV to logistic regression with L2 regularisation from the Sci-Kit-learn library (73) since the non-informative feature’s coefficients were already shrunk to zero by L1 regularisation in the feature selection step. The final regression gave an accuracy of 80% and the MCC or Mathew’s Correlation Coefficient of 0.588. The Receiver Operating Characteristic (ROC) curve revealed an impressive area-under-the-curve (AUC) of 0.844 (**Figure 4A**). The ROC aided in assessing the performance of the model selection and fit. From the confusion matrix in **Figure 4B**, we observed that the model predicted 20 K13 variants as true positives (ART-R), 12 as true negatives (ART-S), with 5 and 3 variants as false positives (ART-S predicted as ART-R) and false negative (ART-R predicted as ART-S), respectively. The model also predicted that the majority of ART-sensitive strains with a P (resistance) < 0.4 and most resistant strains with P (resistance) > 0.5 (**Figure 4C**). The predicted probability distribution histogram in **Figure 4C** showed a near clear separation between sensitive and resistant K13 mutants. To further assess whether the model’s performance was solely due to an indicative signal from the core features and not by change, we performed a permutation test wherein the labels were randomly shuffled and the full nested LOOCV pipeline was re-run on the mislabelled data a thousand times. The permutation test indicated that the accuracies of mislabelled and permutated models peaked between 55% and 60%, compared to the 80% accuracy of the original model that was substantially outside the distribution (**Figure 4D**). Thus, the permutation test yielded a p-value of 0.002, which indicated that the accuracy is highly unlikely to be due to chance. The final deployed model can be found at: “https://github.com/22rajdeep04/k13-ARP.git” under ‘training_scripts/k13_logistic_regression.py.’ The final fit of the model revealed ALIN has the highest coefficient (1.2068) followed by BDE and II (1.1094 and 0.8744 respectively) to calculate the logit(P) values which is used by the sigmoid function to yield the probability of resistance or P (Resistance).

**Figure 4.**
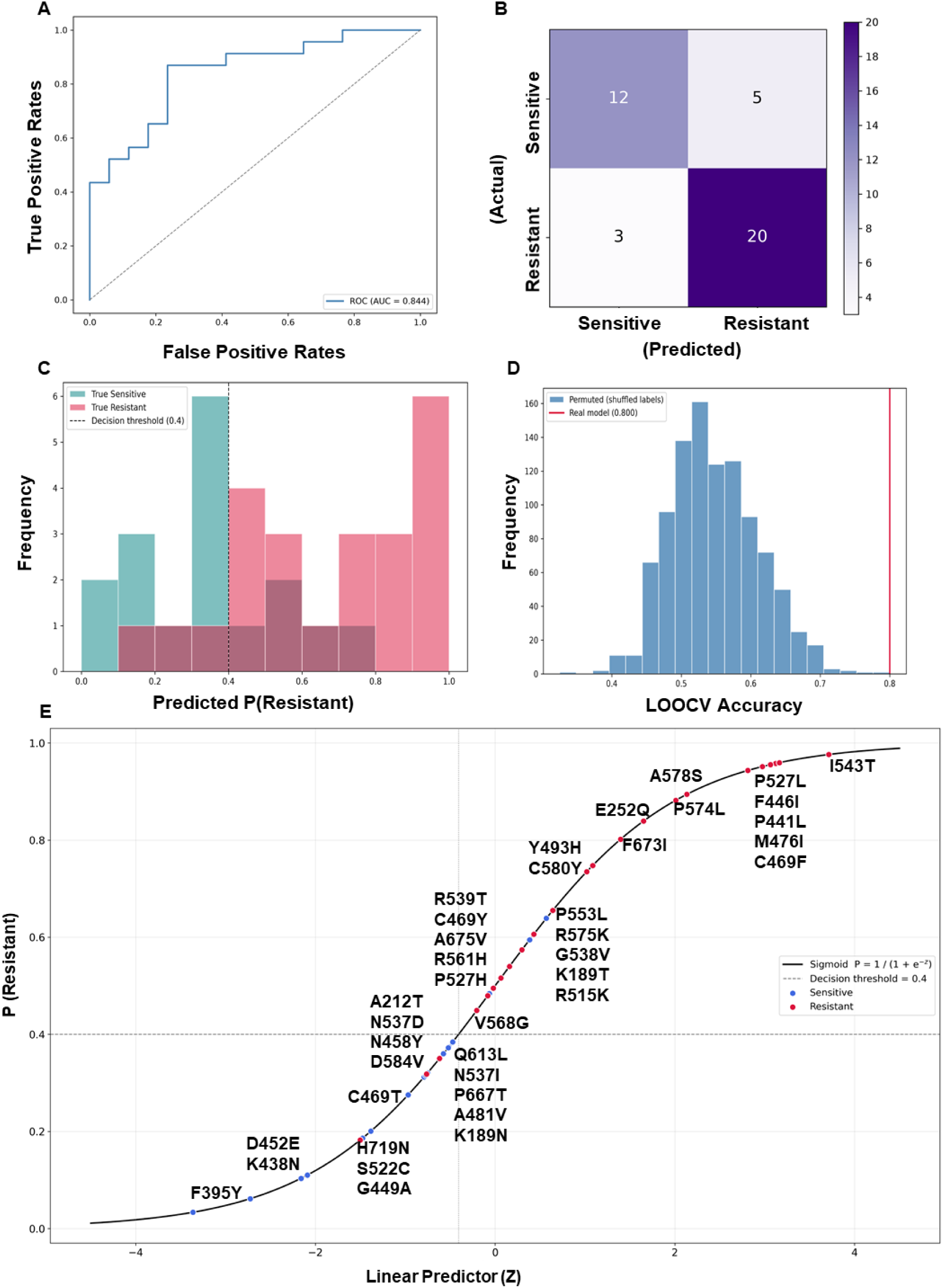
Logistic Regression on selected features showed excellent predictive power. **A.** Receiver Operating Characteristic (ROC) curve of logistic regression model with L2 regularisation and AUC (Area Under Curve) = 0.844, indicating excellent performance of the model. **B.** Confusion matrix (with the decision boundary set to 0.4), revealing the prediction of 20 K13 variants as true positives (ART-R) and 12 as true negatives (ART-S). Only 5 and 3 variants were predicted as false positives (ART-S predicted as ART-R) and false negative (ART-R predicted as ART-S), respectively. **C.** Predicted probability distribution histograms. Red columns represent resistant strains while green columns represent sensitive K13 mutants and wild-type K13. The model distinguishes sensitive from resistant K13 variants by employing a decision boundary at 0.4 (black dotted line). **D.** Histogram showing performance of the model when stress tested with a permutation test. The real model outperformed models trained on mislabelled data in terms of accuracy. **E.** Sigmoid curve representing classifiers predicted probability plotted against the linear predictor, z.

The programme calculated the final output by first transforming the input using the standardisation function where x, µ and σ represented the input value, mean and standard deviation, respectively of the complete data (n=40) in the final refit model. **Table 2** represented the mean and standard deviation of each core feature and the intercept magnitudes of the model.

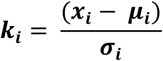

**Table 2.** Standardisation parameters and intercepts of the final deployed models. High mean (µ) and standard deviation (σ) of ALIN and II, as compared to BDE due to the larger range of variations in the ALIN and II scores of the mutants. For any new mutant, the mean and standard deviations were used to calculate the raw score that was then parsed through the model coefficients to calculate the final output. The intercepts of the final deployed models were refit over the entire training dataset of n = 30 for linear regression and n = 40 for logistic regression.

| **Feature** | **μ** | **σ** |
|---|---|---|
| **Logistic Regression** |  |  |
| II | 21.832 | 25.055 |
| ALIN | 32.585 | 32.141 |
| BDE | 1.442 | 1.000 |
| **Linear Regression** |  |  |
| ALIN | 30.022 | 34.153 |
| **Intercepts** |  |  |
| Logistic Regression | 0.2220 |  |
| Linear Regression | 0.4938 |  |

The scaled and standardised input was parsed into the logit function:

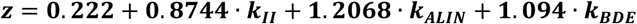

To calculate the probability, the sigmoid function used the previously calculated z as:

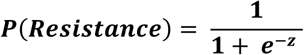

The aforementioned functions yielded a sigmoid curve shown in **Figure 4E**.

### Linear regression model shows weak predictive power for parasite-clearance half-life

After we had generated a model which could efficiently classify K13 mutations into ART-S and ART-R phenotypes, we further explored if our dataset could also quantify the predicted resistance levels as a measure of the PC_1/2_ (Parasite Clearance Half-Life). After an exhaustive feature selection, we used linear regression to build the same model.

Firstly, we referred to composite table (**Table 1**) for the list of mutations and their respective PC_1/2_ (80,81,84). From the same, we also generated the ‘K13_pc.csv’ file (**Supplementary table 3**) with PC_1/2_ value of each mutant instead of the discrete resistant/sensitive labels (**Figure 2**). The linear regression model was trained on only 30 samples due to limitations in validated sources confirming the PC_1/2_ of mutant K13 mutant strains. Next, we used an L1-regularisation based LassoCV to find the most associated features with our target variable in which, we used LassoCV in a Nested-LOOCV to identify the selection rates of each physicochemical parameters considered in this study. Results indicated that only AI was selected in 76% of the folds followed by SH in 3% of the folds, while other features were never selected during the nested LOOCV feature selection process (**Figures 5A-B**). To allow for another independent approach toward the exact feature set, we prepared correlation heatmap to validate the features which are most correlated with the target variable. As shown in **Figure 5C**, we observed a notably higher correlation between AI and PC_1/2_ as compared to other variables. Since only one feature was retained for the final model, a VIF check for multicollinearity was deemed unnecessary. In **Figure 5D**, Q^2^ *versus* k sweep (where k represents the number of features retained for the model), we observed that Q^2^, the cross-validated predictive relevance, modestly peaked between k = 1-2. However, when we selected only AI for the final model, the cross validated LOOCV Q^2^ value reached 0.110. With both AI and SH selection, Q^2^ value showed an infinitesimal increase to 0.121, which was only a 10% increment. Considering the modicum selection rate of SH (3%) compared to that of AI (76%), and the inability of the model (with both SH and AI) to substantially improve the predictive power, we speculated that the marginal gain in Q^2^ was unable to justify retaining SH as a core feature, and a simpler model that stably converges was a parsimonious but judicious and astute choice, since SH might have introduced noise rather than genuine predictive power.

**Figure 5.**
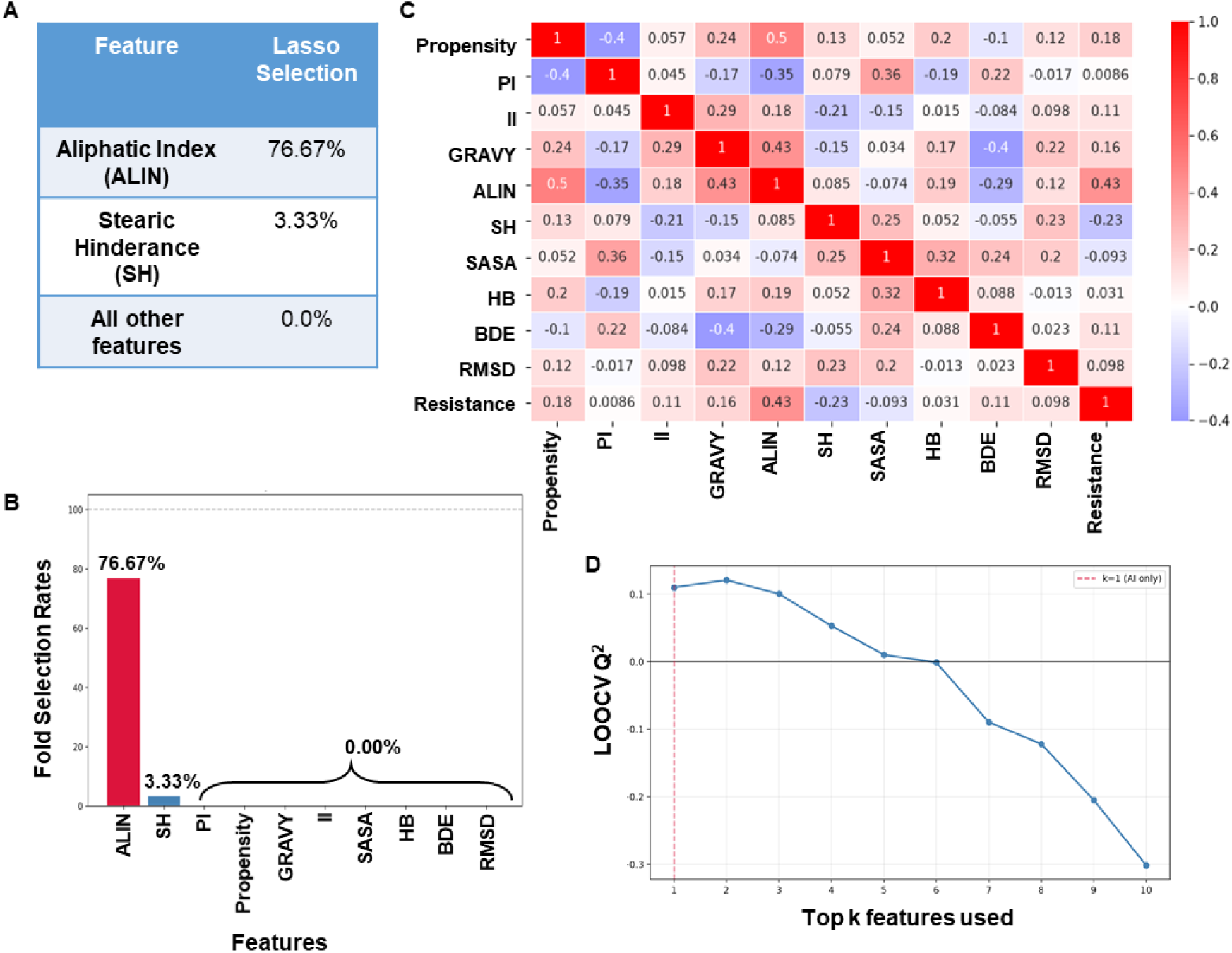
Exhaustive feature selection criteria for the identification of collinearity of features with the parasite clearance half-life. **A-B.** Lasso selection table showing high selection frequency for ALIN but not any other feature, thereby indicating low predictive power of linear regression on the training dataset. **C.** Correlation heatmap indicating high correlation, |r| = 0.43, of ALIN with the parasite clearance half-life, when compared to the rest of the parameters. **D.** Q^2^ versus k curve showing peak performance at k = 2 (0.121), although only at 10% higher than at k = 1 (0.11), which was indifferentiable from noise. Hence ALIN was chosen as the only core feature.

Since the PC_1/2_ values represented a ratio of parasite clearance rate in K13 mutants as compared to that of the wild-type K13, it introduced a rather log-linear relationship with the features. Hence, transforming the PC_1/2_ as the target variable empirically improved the Q^2^ score from 0.066 to 0.110. The in-sample R^2^ value of 0.217, which was considerably higher than the cross-validated Q^2^. Once the model was trained, and cross validated, we plotted the predicted *versus* true value plot (**Figure 6A**), which showed a rather cloud-like orientation. The residual plot in **Figure 6B** also revealed no apparent pattern, indicating a lack of ALIN’s ability to stably predict and capture the variations in PC_1/2_. We then trained a separate unregularized OLS (ordinary least square) regression to assess the statistical significance between the log (half-life) and ALIN scores of the mutants. In **Figure 6C**, we noted that the p-value for the regression was 0.0095 and the slope of the graph was 0.0049. We also observed 95% confidence interval around the regression line, to factor for the ambiguity in the position of the regression line. The 95% confidence interval widened at the edges and narrowed down at the centre, which was consistent with the input data since there were not many samples at either extreme and plenty within the centre. Similar to the logistic regression, we performed a permutation test to confirm that the observation was not by chance. Unlike the logistic regression, which gave a p-value of 0.002, the linear regression only gave a p-value of 0.013 (**Figure 6D**), which also revealed that there were a few mislabelled models that returned a Q^2^ equal to or more than that of the real model. The final refit model has been made publicly available and can be found at: “https://github.com/22rajdeep04/k13-ARP.git” under ‘training_scripts/k13_linear_regression.py.’ The model only requires ALIN as the input and can predict the parasite clearance half-life of the input K13 variant.

**Figure 6.**
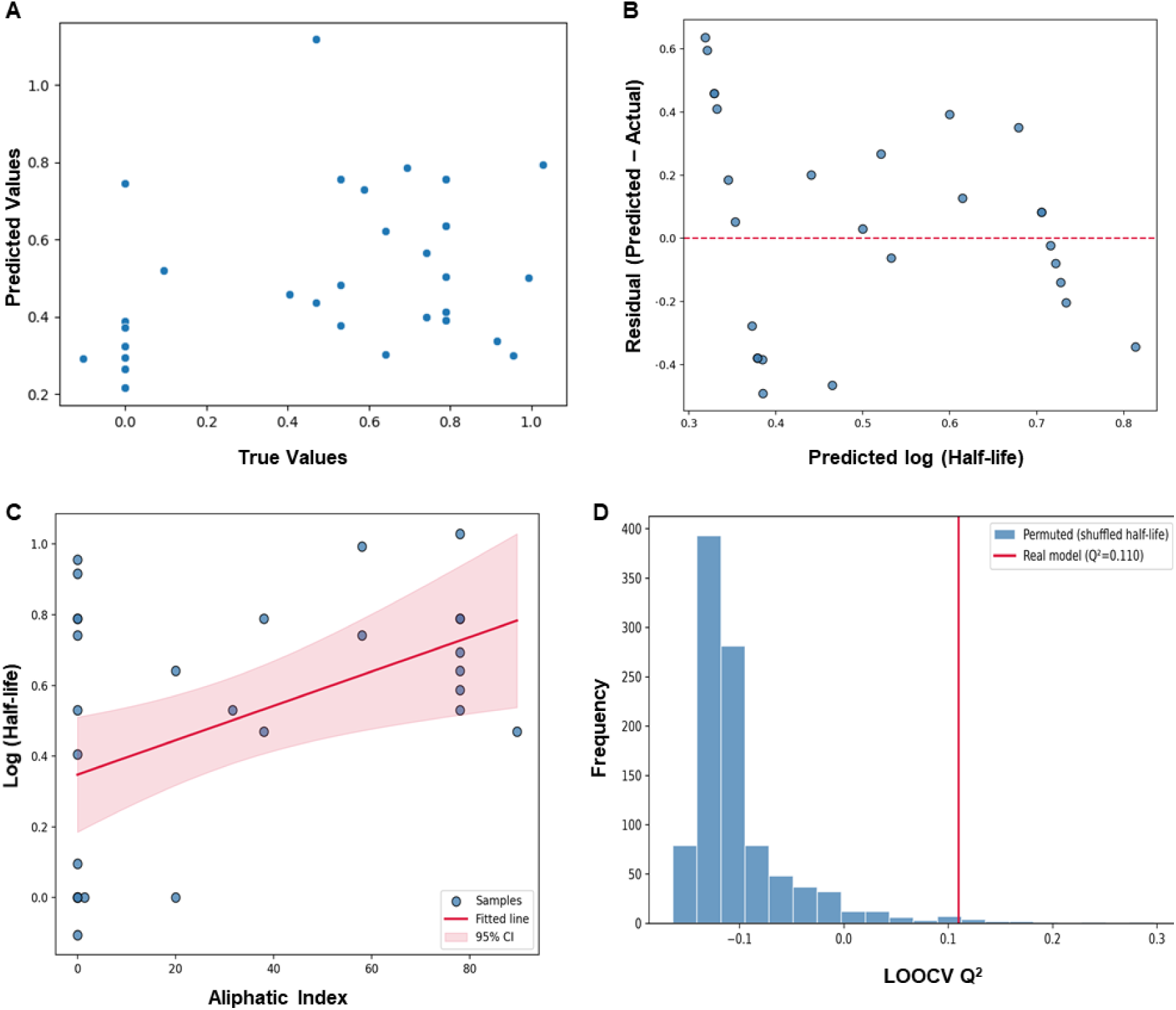
Poor performance of linear regression on the harvested dataset indicating the involvement of more complex set of features for PC_1/2_ values. **A.** Graph showing cloud like arrangement between predicted and true values and indicative of lack of effective collinearity between the core feature ALIN and PC_1/2_. **B.** Similar appearance (as in A) and the lack of an apparent pattern such as a parabola in the residual plot. **C.** Graph showing 95% confidence interval (red line) around the exceptionally broad regression line and indicative of a lack of confidence in the regression in the model consistent with the low Q^2^ = 0.11. **D.** Histogram showing outperformance of most models trained on mislabelled data when stress tested with a permutation test and reflecting that the model’s scores were not accidental.

The model standardised the input variable using z score normalisation where the difference of the input and mean µ (30.022) was divided by the standard deviation of the data, σ (34.153).

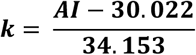

Since the relation between PC_1/2_ follows a log-linear relation, the final output could be exponentiated as:

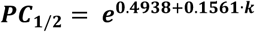

## DISCUSSION

Current challenges in assembling the metadata generated from malarial epidemiological surveillance, population migration and climate variability often lead to questionable data quality, limitations in infrastructural facilities and conformant to ethical guidelines. Undoubtedly, the vast and heterogeneous datasets require advanced analyses to extract relevant insights (85–87). On deliberation, we sought AI-based computational methods, including machine-and deep-learning tools to enable *in silico*-based algorithms which mimic human intellectual abilities by learning from data, identifying patterns, and making predictions. AI-based predictions also offered an attractive opportunity to elucidate the molecular mechanisms underlying resistance to the frontline antimalarial ART derivatives (implemented as ACT), in conjunction with previously published *in vivo* or *in vitro* transcriptomic traits (16,50,88–90).

Till date, only a few prior studies have evaluated the complex relationship between mutation(s) in individual genes and their expression levels (91,92), although a systematic evaluation of K13, the leading molecular marker for ART-resistance, has not yet been attempted. More than 200 non-synonymous mutations or Single Nucleotide Polymorphisms (SNPs) have been globally identified and mapped within the *P. falciparum* K13 sequence (32,56), primarily localized in the highly conserved C-terminal KRP domain of the protein (23). Around 13 specific mutations in K13 have been strictly validated and statistically proven to correlate with delayed parasite clearance in patients and display survival advantages under laboratory conditions; and roughly 9 mutations have been officially classified as “candidate” markers, suspected of causing resistance due to field observations but still lacking definitive, isolated laboratory proof (1,56,93). Overall, this study was designed to harness AI-based algorithms and develop a screening platform for a representative set of K13 variants, which could then be rapidly translatable onto a robust and time-efficient predictor for broad ART-responsive phenotype. The outcome demonstrated that a parsimonious set of 10 physicochemical and structural descriptors, derived from sequence and AlphaFold-predicted structures, was capable of predicting K13-mediated ART-R with 80% accuracy. The logistic regression model robustly classified ART-R *versus* ART-S mutants, although it incorrectly classified eight of the 40 held-out samples. Notably, the three false-negative resistant mutants (N458Y, N537I, G449A) exhibited aliphatic indices near the lower threshold of the resistant class, suggesting a continuous resistance gradient rather than a binary switch. Our findings aligned with recent reports implicating that R561H and A675V displayed variable clearance delays across African settings (94). Our model’s high accuracy (80%) and AUC (0.844) also compared favourably with recent machine learning (ML)-based approaches for drug resistance prediction in other pathogens, which reported pooled AUCs of 0.8-0.9 (95–97). While our sample size (n=40) was modest, yet rigorous cross-validation and permutation testing supported the generalizability of the learned patterns. Importantly, the models relied solely on sequence-and/or structure-derived features, obviating the need for expensive phenotypic assays or transcriptomic profiling. In addition, it offered the potential for deployment as a rapid screening tool for emerging K13 mutations in resource-limited settings.

Our descriptor analysis yielded several mechanistic insights for predicting K13-mediated ART-responsiveness. The propensity calculations delineated the putative ART-S mutations in K13 from the ART-R ones, and revealed only minor deviations in the former variants as compared to the latter, with major deviations in the K13 mutant which has been considered as highly ART resistant [such as R539T, (16,88)], **Figure 7A**). We recently reported that the relative binding affinity of heme for the K13 protein played a key role in the activation of pro-drug ART into metabolically active heme-ART adducts, carbon-centered C4 radicals and toxic reactive oxygen species (ROS) (98). Thus, any physicochemical parameter that would potentially disfavour heme stability and its binding to K13 was further evaluated. As previously reported, heme binding favour neutral to basic pH conditions depending upon the ionization state of the axial coordinating amino acids and the structural stability of the host protein. Under acidic conditions, the binding affinity drops significantly, often leading to heme dissociation (99). Our results (**Supplementary table 1**) confirmed that the well-known (and previously-validated) ART-R K13 mutant (R539T) reduced the local pI, increased the net negative charge within the heme-binding pocket and dis-favoured heme binding at physiological pH as compared to the ART-sensitive wild-type K13, which perfectly aligned with the reduced ART-activation model, wherein diminished heme-K13 interactions decreased ART activation (98). In contrast, the K13 F446I mutant showed lower deviation from the wild-type protein (**Figure 7B)**. Incorporation of GRAVY score further showed similar yet distinct deviation from wild-type K13, where the previously-validated highly resistant strains (based on their RSA index P574L, R539T, C580Y etc) deviated further from WT than the moderately resistant strains with K13 mutations (F446I, K189N), shown in **Figure 7C**.

**Figure 7.**
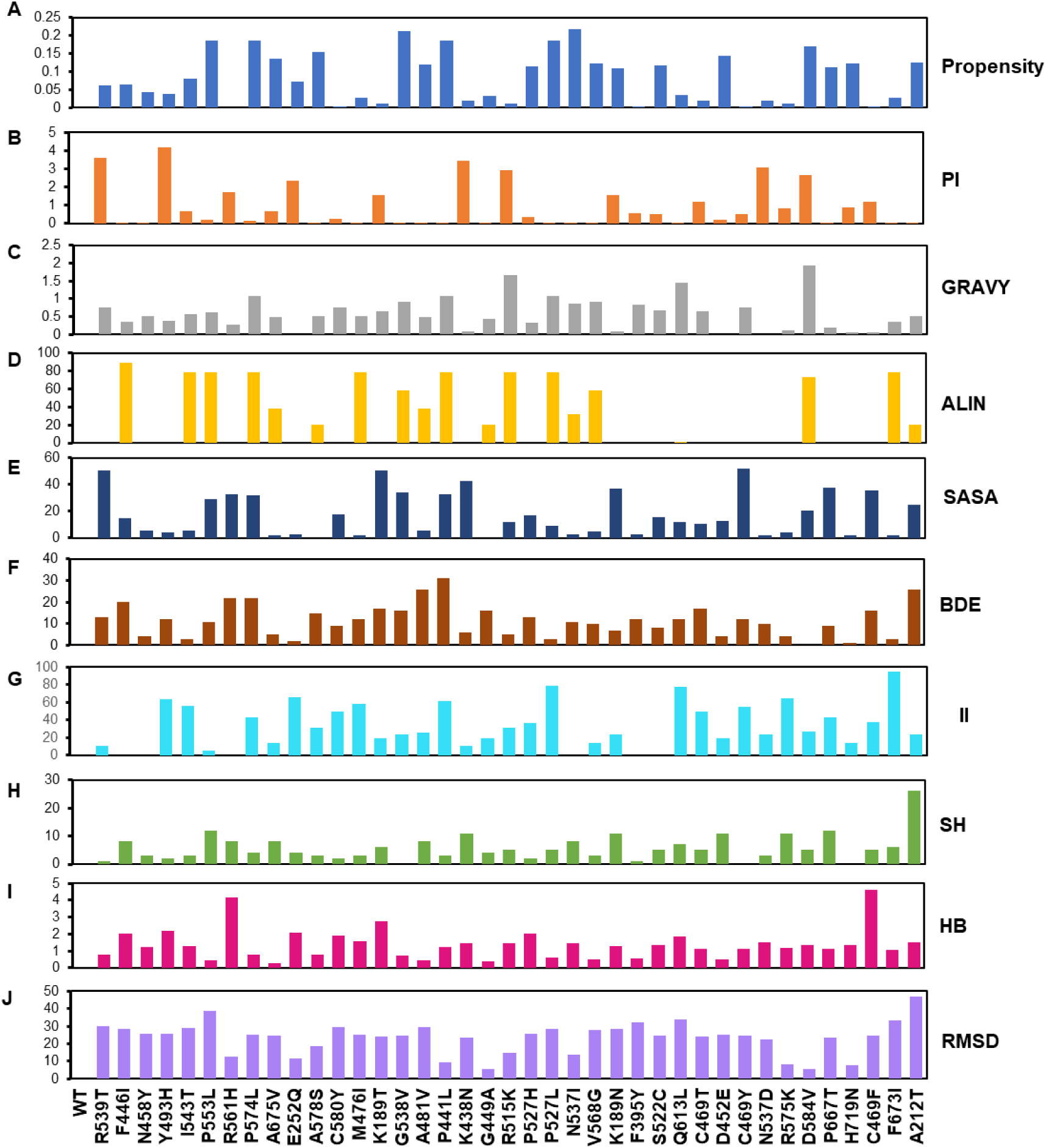
Comprehensive and exhaustive representation of data acquired from various tools and servers for individual parameters/features. **A.** Propensity distribution data showing that clinically identified ART-R mutant with K13 A675V displayed higher deviation in the calculated propensity as compared the wild type (WT) protein than the ART-S K13 mutants (*e.g.*, K189T. **B.** PI data indicating that resistant strains (R539T) displayed values with high deviation from the WT K13 than sensitive mutants (*e.g.*, F446I). **C.** GRAVY score measuring the hydropathicity of the local environment. **D.** ALIN measurement (of the aliphaticity of the local environment and contribution to the protein’s ability to efficiently bind the heme macrocycle) showing deviation of validated ART-R K13 mutants from WT were less than the ART-S K13 mutants. **E.** SASA parameter showing similar pattern as propensity. SASA is directly indicative of how buried a residue may be in the three-dimensional conformation of the protein, with higher perturbation leading to high resistance downstream. **F.** BDE measurement of thermodynamic change at a dipeptide level and showing similar pattern as ALIN. **G.** II data showing that although it lacked of an apparent pattern between resistance phenotypes, yet it was crucial to thermodynamic stability of the protein. **H.** SH or clashes between sidechains of amino acid residues in the sequence of the protein leading to packing instability. Too many clashes would likely render the protein non-functional. K13 mutations conferring ART-R deviated conservatively than ART-S K13 mutants **I.** HB profile showing conservative deviation for validated resistant mutations in K13 as compared to sensitive mutants. An increase in HB correlated to much tighter packing, leading to loss of certain functions in K13. **J.** Structural deviation, as represented by RMSD showing values for ART-R K13 mutants than the ART-S variants.

Membrane-association of K13 protein has been previously reported (100). Both endogenous K13 and green fluorescent protein (GFP)-tagged K13 proteins were found to be preferentially localized at membranous compartments proximal to the parasite food vacuole (FV) and cytostomal structures near the periphery of the parasite (101,102). Thus, any alteration in local hydropathicity would theoretically mislocalize the protein, leading to inconsistent activation of ART or the formation of heme-ART adducts, thereby allowing resistant mutant(s) to survive the ART exposure (103,104). Indeed, conditional mislocalization of K13 using the diCre-based approach resulted in developmental arrest of the intraerythrocytic parasite at the late ring stage where the ART-resistance clinically manifested (102,103). Because the heme macrocycle is hydrophobic, it binds preferentially to hydrophobic protein pockets (105,106). The ALIN analyses of K13 mutants previously-validated as ART-R (such as A675V and R539T) revealed smaller deviations from the wild-type K13 protein. In contrast, the K13 mutant F446I deviated more strongly (**Figure 7D**), which suggested that a definitive phenotype for ART-resistance involved a composite from multiple physicochemical parameters which resided in a ‘Goldilocks zone’, *i.e.*, deviating sufficiently to confer resistance while preserving parasite viability. Given that ART activation was more efficient when heme was bound to K13 (98), thermodynamic destabilization of the protein would reduce heme-ART interactions. The SASA profile also exhibited pattern analogous to pI, wherein the highly resistant K13 mutations showed larger absolute deviations from the wild-type K13, implying that the respective mutations (R539T, P574L) either buried or exposed more amino acid residues, which likely increased the solvent exposure to free cytosolic heme or ART and thereby reduced ART activation (**Figure 7E**). Similarly, reduction in local BDE at the site of mutation implied weakened intramolecular bonding, thereby contributing directly to the fold-destabilization pathway (107,108). Since evolution of K13-mediated ART-resistance most likely involved many independent constraints, our investigation into the global physicochemical parameters and structural changes which could potentially destabilize the K13 protein or its heme binding ability of K13 directly captured the thermodynamic cost of mutation (**Figure 7F**). II deviation from WT shows no discernible pattern, although validated resistant strains such as C580Y and I539T deviated much further than clinically sensitive strains like F446I, other resistant strains such as R539T and A675V do not show a high deviation from WT (**Figure 7G**). This finding is coherent with the nature of the index, since it is calculated at a dipeptide scale, and therefore it only captures the sequence-limited thermodynamic instability locally rather than structural or functional consequences. Our data exploration revealed that the deviations SH was consistent with the pattern shown by ALIN, where the ART-resistant K13 mutants like R359T and C580Y showed lesser deviation than predicted ART-sensitive mutants like F446I and S522C (**Figure 7H**).

AlphaFold generated 3D structures have previously been shown to conservatively change the number of hydrogen bonds (HB) in serpin mutants as compared to the wild-type protein (109). In context to K13, prominent ART-resistant mutants (such as C580Y, R539T, and Y493H) have been proposed to alter the protein structure just enough to cause reduction in protein function or abundance, thereby limiting hemoglobin uptake and avoiding ART activation. However, ≥ 90% of the original intra-chain hydrogen bonds still remained in the ART-R K13 mutants (102,110,111). Ultimately, AlphaFold predictions have confirmed that successful ART-R mutants strike a perfect evolutionary balance: while they subtly adjust protein dynamics and binding affinity (such as lowering heme-binding capacity), the core scaffold of hydrogen bonds (**Figure 7I**) remained highly conserved to maintain vital structural integrity (21,98).

RMSD deviation measured the structural deviation of the K13 mutants from the wild-type protein (**Figure 7J**). Mutations associated with larger RMSD were more likely to perturb the local packing of the protein’s 3D-conformation, thereby leading to structural instability. We incorporated the RMSD tool as a proxy for the structural perturbations rather than a predictor of loss of function. The KRP domain formed a closed and densely packed structure with six β-sheets making each of the six propellers. The dense arrangement was only possible if the side chains of individual amino acids did not clash with their adjacent amino acid’s side chains (111)

Upon training and fine tuning, the logistic regression model achieved an accuracy of 80%, which meant that the model was able to correctly classify 80% of the held-out samples using only physicochemical and structural descriptors. In total, it only mis-classified eight mutants, namely N458Y, N537I, G449A, K189T, K189N, Q613L, C469Y, R575K, where the first three have been categorized as ART-R but were classified as ART-S by the model (False Negative; FN), and the latter five were classified as ART-R but were indeed ART-S (False Positive; FP). A common pattern emerged after further investigation of the misclassified mutants. The false negatives shared a low ALIN score relative to the correctly classified ART-resistant group, which was consistent with the ART resistance phenotype of candidate marker N537I and the experimentally validated G449A, which was deemed closer to the lower threshold of ART-resistance. In the case of K189T, the FP mis-classification was primarily due to II and BDE, since its ALIN value was exactly 0. Although, based on ALIN alone it should have been classified as ART-S. Since ALIN has the highest coefficient on the final refit model, it contributes the most towards the resistant classification of the mutant by the model. The other features are unable to compensate for the complete loss in the contribution of ALIN, which results in false predictions such as that of N537I and G449A.

However, with the small sample size of n = 40, we had to deliberately ensure that the model could confidently generalise the output, instead of learning from the quirks of the small dataset. The permutation test revealed that the model was able to outperform any mislabelled dataset over 1000 iterations. Since our model was capable to predict the resistance index of the mutants, we further explored if the dataset could also reveal patterns in PC_1/2_ values. However, on training a separate linear regression model with the PC_1/2_ values as the only target variable, we observed that ALIN alone showed weak mathematical correlation with continuous predictions. The model achieved a Q^2^ of 0.110, which indicated a weak but potentially real signal. The Q^2^ *versus* k sweep revealed that Q^2^ declined beyond k = 1-2. Convening a permutation test discovered that the model underperformed as compared to a few models trained on mislabelled data, thereby indicating that ALIN was unable to predict PC_1/2_ and the residual diagnostics showed no systematic pattern. The linear regression model’s poor performance (Q² = 0.110) further highlighted the complexity of predicting parasite-clearance kinetics from sequence features alone. Integration of host factors (*e.g.*, immunity, drug metabolism), parasite transcriptomics, or pharmacokinetic data may be necessary for accurate quantitative predictions (112–114). To generate insights into novel and emerging mutant strains, we predicted the P(Resistance) score of the mutant and classified them accordingly. Out of the 18 unclassified mutant strain, the model predicted 6 as sensitive and 12 as resistant. **Supplementary table 4**, and **Figure 8A** shows the P(Resistant) score of the mutants, allowing new mutants to be classified and labelled as drug-resistant even before they emerge or cause a large-scale epidemic, taking away millions of innocent lives, in a time-efficient manner without the need to perform laborious lab adaptations or RSA in impoverished endemic regions across the world. The sigmoid curve of P (Resistance) plotted against the linear predictor logit; z, in **Figure 8B** showed that the novel mutants fit onto the original refit of the classifier. Since the linear regression model only ran on the ALIN input of the mutant sequence, the predicted PC_1/2_ incorporated only 3 values corresponding to the three possible ALIN differences observed during data collection.

**Figure 8.**
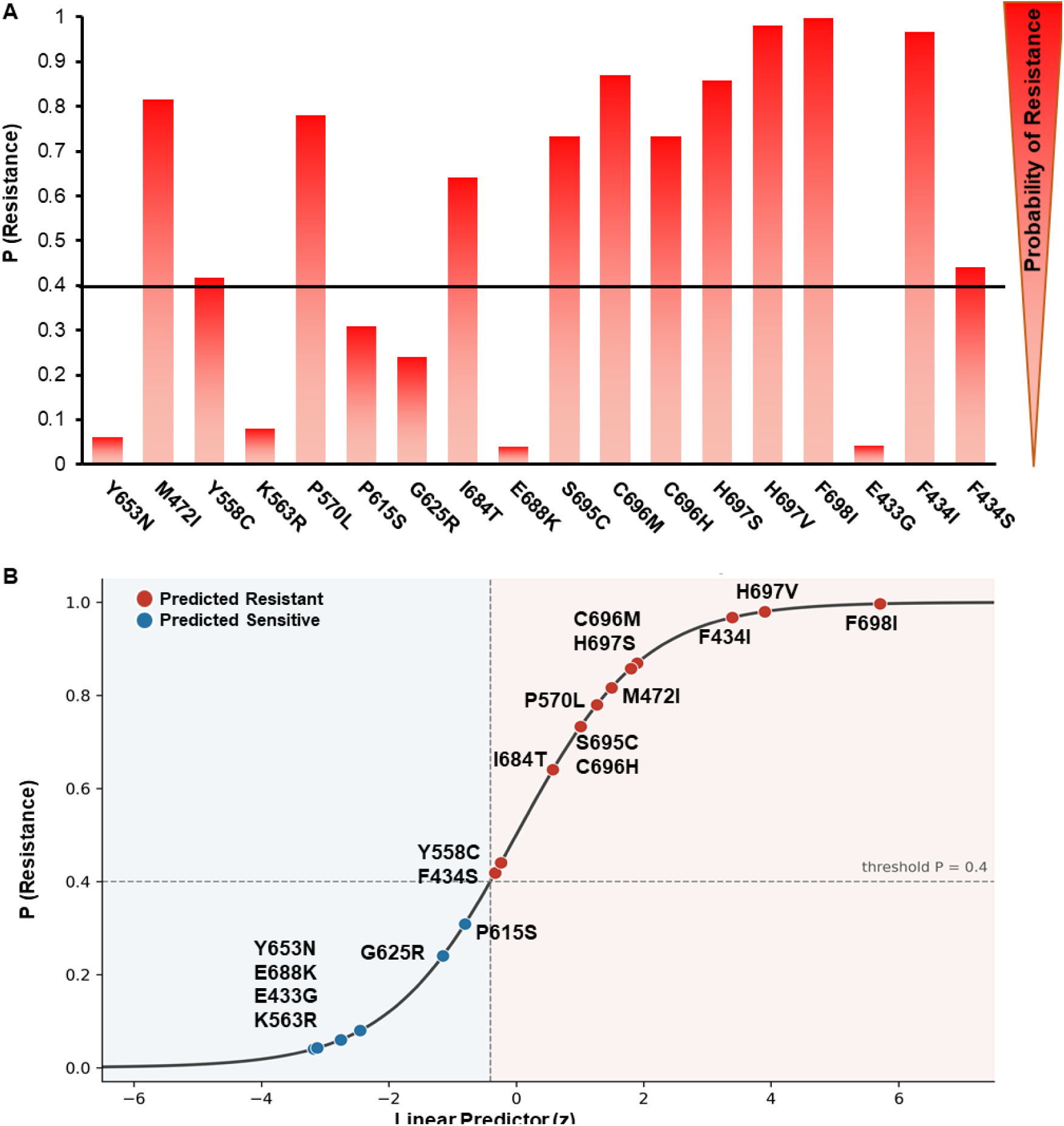
Predictions for recently reported and previously uncharacterised mutations in K13. **A.** Bar graph showing that P (Resistance) of the 6/18 recently reported K13 mutants is below the threshold (0.4). For 10/18 mutants in K13, it was significantly higher than the threshold. Only 2 K13 mutants (Y558C and F434S) were slightly higher than the threshold. **B.** Sigmoid curve displaying the linear predictor and output P (Resistance) for K13 mutants predicted as resistant or sensitive. P615S, G625R, Y653N, E688K, E433G and K563R were predicted as sensitive to ART. Although the model predicted Y558C and F434S as resistant, their confidence was significantly lower than any other predicted resistant mutant, thereby indicating a low confidence call.

Although the logistic model converged and generalised to great extent, it left opportunities for further developments. The primary limitation of this study was the small sample size (n=40 for model training and validation), which constrains statistical power and generalizability. While cross-validation and permutation testing mitigate overfitting, larger datasets encompassing geographically diverse mutations (particularly from Africa) would strengthen model robustness, additionally, collapsing low, medium, and high resistance into a binary class obscures quantitative differences in ART-resistance severity; future work should explore multi-class or ordinal regression models with larger datasets. Nonetheless, deployment of this ML framework could accelerate molecular surveillance of ART-R, particularly in Africa, where phenotypic assays are scarce and genotypic data are fragmented. By rapidly classifying emerging K13 mutations as resistant or sensitive, public health programs could prioritize high-risk variants for further investigation and inform ACT policy decisions. Integration with existing surveillance platforms line MalariaGEN [https://www.malariagen.net; (115,116)].

## Supporting information

Supplemental Taable 2

## AUTHOR CONTRIBUTIONS

Conceptualization Ideas: R.D., S.D. and S.B.; Methodology: R.D.; Validation: R.D., Investigation: S.D., S.B.; Resources: S.D. and S.B., Writing and edit: R.D., S.B., S.D.; Supervision: S.D. and S.B.; Project Administration: S.B.

## ACKNOWLEDGEMENTS

This work was supported by research grants from the Indian Council of Medical Research (ICMR), (58/25/2020/PHA/BMS) and Indian Ministry of Human Resource Development (MHRD) Scheme for Translational and Advanced Research in Sciences (STARS) (MoE-STARS/STARS-2/2023-0111) from the Government of India (SB). SD is a recipient of Department of Biotechnology Research Associateship (DBTRA/2023 24/N/JNU/118) from the Government of India.

## GLOSSARY AND ACRONYMS

Local Parameters: refers to those physicochemical or structural parameters that only affect the region of the protein surrounding the mutations and not the complete protein structure.
Global Parameters: refers to the parameters that were observed to be concerned with the entire protein’s physiochemistry or more often but not exclusively the structure.
WT: Wild type
GRAVY: Grand Average of Hydropathicity
PI: Isoelectric Point
SASA: Solvent Accessible Surface Area
HB: Hydrogen Bonds
SH: Steric Hindrance
II: Instability Index
ALIN: Aliphatic Index
BDE: Bond Dissociation Energy
RMSD: Root Mean Square Deviation
K13: *Plasmodium falciparum* protein Kelch13

